# Variations in cow milk and teat skin microbiota across the lactation cycle with intra-mammary cephalosporin use at dry-off

**DOI:** 10.1101/2025.11.19.689190

**Authors:** Wannes Van Beeck, Mateus Lemos, Ashley M. Niesen, Peter Finnegan, Taylor M. Shih, Anhha Ho, Heidi A. Rossow, Maria L. Marco

## Abstract

Cephalosporins and other broad-spectrum antibiotics are frequently administered prophylactically into the udder when dairy cows end their lactation cycle, termed dry-off, to reduce mastitis risk. However, the use of antibiotics on cows that do not have signs of infection may result in the selection of microbes resistant to antibiotics and negatively alter the udder microbiome composition. In this study, the effects of intramammary cephalosporin therapy with either Cephapirin (CB) or Ceftiofur (CH) on milk and teat skin microbiota were examined for three dairies in California. Bacterial composition was measured for cows with low somatic cell counts (SCC,<100,000 cells /mL) and high, subclinical SCC (>200,000 cells/mL). Samples were collected at dry-off (before treatment), seven days later, and 55-75 Days in Milk (DIM) in the next lactation cycle. The milk and skin microbiota were largely separated based on dairy (milk: R^2^ = 6.22, skin: R^2^ = 7.56) and day of sampling (milk: R^2^ = 4.74 and skin: R^2^ = 3.77). CB or CH use was associated with a small but significant impact on the milk microbiota beta-diversity (Bray-Curtis, p =0.003, R^2^= 1.4%) but no effect was observed on the skin. At one dairy (Dairy 3), milk from cows receiving CB and CH had a reduction in proportions of *Staphylococcaceae* at 55-75DIM compared to untreated cows. Overall, antibiotic use did not result in large significant changes to bacterial diversity in milk or on the teat skin, and instead the microbiota at those sites mainly differed between the time and location of sampling.

**Importance:** The use of antibiotics in agriculture is under increasing scrutiny due to the rising spread of antimicrobial resistant bacteria. Our study showed that common preventative antibiotic intramammary treatment of cows with cephalosporins at the end their lactation (dry-off) had minimal effects on the milk and teat skin microbiota on asymptomatic cows with high somatic cell counts.

## Introduction

Intramammary treatment with broad-spectrum antibiotics is frequently used as a preventive measure for dairy cows at the end their lactation cycle (also called dry-off) to reduce the risk of infection and inflammation (1, 2). Up to 80% of dairies across the United States have adopted this prophylactic strategy (3). However, prophylactic, intramammary antibiotic use for cows that do not have signs of infection may pose a risk of increasing the spread of antimicrobial resistant microorganisms and changes to the mammary microbiome resulting in increased mastitis susceptibility in subsequent lactation cycles (4). Additionally, parenterally administered antibiotics have been detected in bovine milk up to nine days after antibiotic treatment (5). Therefore, the necessity and impact of routine antibiotic use at dry-off as preventive measure has come under question.

Bovine milk from actively lactating cows contains diverse microorganisms belonging to multiple phyla including the Bacillota (formerly known as Firmicutes), Pseudomonadota (Proteobacteria), Actinomycetota (Actinobacteria), and Bacteroidota (Bacteroides) (6). In Holstein cows, *Pseudomonas, Streptococcus, Acinetobacter* and *Bacillus* were found to be frequently occurring genera in raw milk (7). The composition of milk microbiota changes during lactation, such as with notable increases in Actinomycetota late in lactation (8). In another study, proportions of *Staphylococcus* were higher late in lactation, whereas *Bacteroides* declined over time (9). The milk microbiota can also carry pathogens, such as with cows with (sub)clinical infections (14, 15). The cow teat skin harbors similar taxa as found in milk and includes species of *Streptococcus*, *Actinetobacter, Staphylococcus, Corynebacterium,* and *Turicibacter* (11). Teat skin microbiota may impact milk quality since microorganisms can be carried over throughout lactation (11). Although the importance of commensal microorganisms in milk and on the teat skin is generally not known, previous studies have associated some genera, such as *Lactobacillus* and *Paenibacillus,* with clinically healthy cows, without infections, and thus may contribute beneficially to the overall health of the cow (12, 13).

Subclinical bovine mastitis is one of the most common issues at dairy farms and is indicated by a decline in milk production, high somatic cell counts (SCC) (>200,000 cells/mL), but no clear sign of teat or udder inflammation (14, 15). A high SCC results when there is an influx of somatic cells from the blood into the mammary gland (16) and suggests damage to milk-producing epithelial cells and underlying inflammation (17). Dairy cows are more prone to udder infection and high SCC during the dry-off period because of factors including slow mammary gland involution, late teat canal close-up, and increased milk retention (18–21). Cows with subclinical mastitis may eventually progress to clinical mastitis or become chronic carriers if left untreated (22). Because of these risks, cows in the United States frequently receive intramammary injections of either Cephapirin (CB), a first-generation cephalosporin (23), or Ceftiofur (CH), a third-generation cephalosporin (24), at dry-off irrespective of SCC status. Both cephalosporins are characterized by their broad activity against both Gram-positive bacteria (such as *Staphylococcus*) and Gram-negative bacteria (such as *Escherichia coli*, *Klebsiella* spp., and *Enterobacter* sp.) (25). Thus, while cephalosporins can inactivate mastitis-causing pathogens, the antibiotics may also harm beneficial commensal microbes in milk or on the teat skin, resulting in acute drug-induced microbiota perturbation.

Herein, we characterized the microbiota in freshly expelled bovine milk and on the teat skin from which the milk was obtained at dry-off (baseline), seven days later, and after 55 to 75 days in milk (DIM) in the next lactation (**Figure 1**) at three dairies in California. At each dairy, groups of cows (10 per group) with high SCC (>200,000 cells / mL) at baseline received an intramammary prophylactic administration of CB or CH or were not treated with antibiotics (high SCC controls). Cows with low SCC (<200,000 cells / mL; 10 per group, low SCC controls) were also included for comparison. The strength of the study lies in its multifactorial approach including the sampling of milk and skin microbiota from cows at different dairies over time with the aims to evaluate milk composition as well as the long-term effects on the bovine udder microbiota in response to prophylactic antibiotic (CB or CH) treatment.

**Figure 1:**
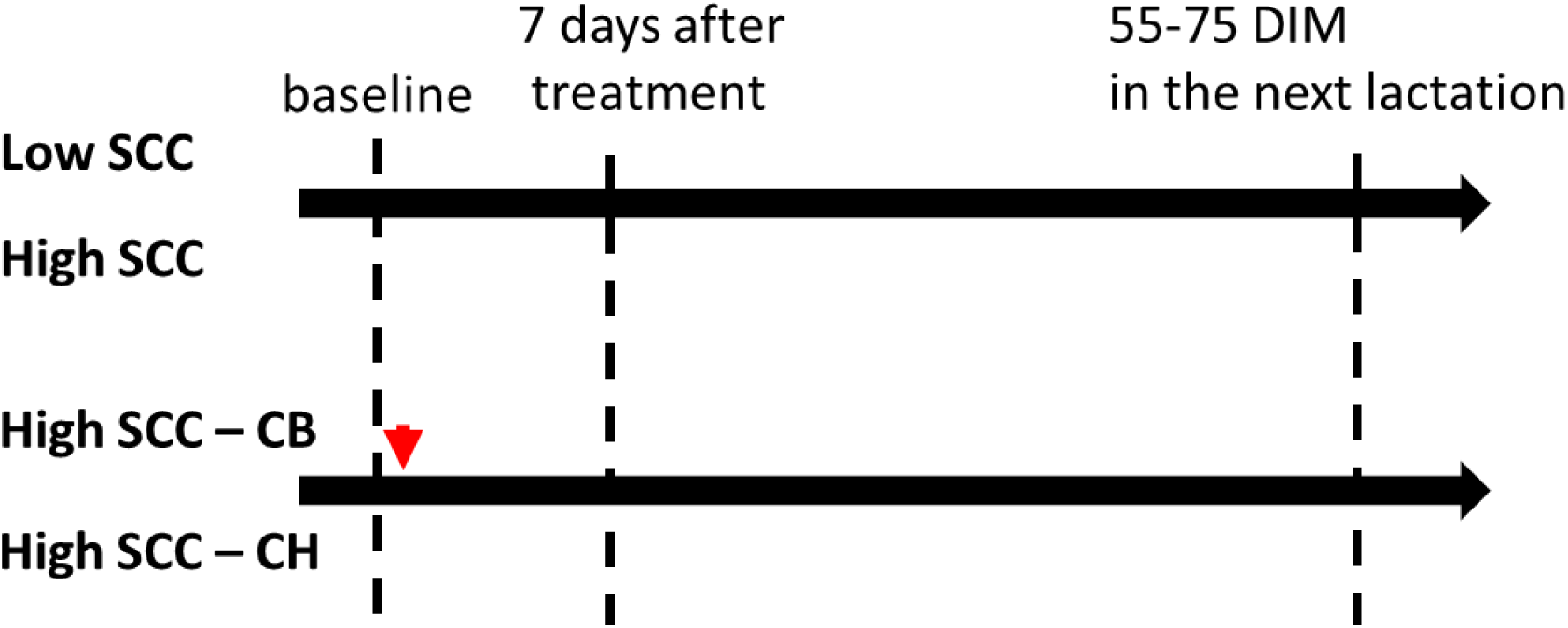
Study design. Milk and teat skin swab samples were collected from three dairies in California, encompassing three timepoints (baseline (dry-off), 7 days later, and 55-75 Days in Milk (DIM) in the next lactation cycle) and four treatment groups (low and high Somatic Cell Count (SCC) controls and high SCC cows given either cephapirin (CB) or ceftiofur (CH) after baseline collection. The red arrow indicates antibiotic treatment.

## Material and methods

## Experimental design

A total of 372 milk and teat skin swab samples were collected for analysis from three dairies in California, encompassing three timepoints (baseline (dry-off), 7 days later, and 55 to 75 DIM in the next lactation) and four treatment groups (low and high SCC controls and cows given either CB or CH) (**Figure 1**). Cows were enrolled in the study if they were greater than 210 days pregnant or if they produced < 20 kg of milk per day. At baseline, milk samples were obtained from each quarter and SCC quantified by the Tulare county Dairy Herd Improvement Association using a Fourier Transform Spectrometer 600 Combi System (Combi; Bentley Instruments, Chaska, MN, USA). Cows were placed in the low SCC group if the SCC were < 100,000 cells/mL in all four quarters. If SCC were > 200,000 cells/mL in one quarter, cows were placed in the high SCC group.

Once cows were grouped by SCC, they were assigned to one of four treatments (n = 10 cows per group): low SCC Controls ; high SCC Controls; and high SCC Cephapirin Benzathine (CB) (300mg ToMorrow; Boehringer Ingelheim Vetmedica Inc., St. Joseph, MO); high SCC Ceftiofur Hydrochloride (CH) (500mg Spectramast DC; Zoetis Inc., Kalamazoo, MI). All treatments were administered by dairy employees. After baseline sample collection and prior to infusion of the cephalosporins, the teats were dipped with an antiseptic (pre-dip), dried with a clean towel, and milk was expelled. The CB and CH treatments were then provided by gently inserting a nozzle containing CB or CH into the teat canal. Cephalosporin was infused (300 mg CB and 500 mg CH) and the teat was briefly massaged. Control treatments received no cephalosporin antibiotics or placebo. Notably, although at least 10 cows were enrolled in each group, some cows were removed over the course of the study because they were no longer available (sold, died, or developed mastitis) (**Figure S1**).

### Teat skin swab and milk collection

The quarter (teat) with the highest SCC per cow was selected for milk and teat swab sample collection and analysis. Cow teats were swabbed with a sterile 2x2 gauze moistened with sterile NaCl (.9% w/v)-Tween 80 (1% v/v) stored in a 50 ml tube. Sterile hemostats were used to remove the gauze from tube and swab the teats. For milk collection, the external teat surface was wiped with gauze soaked in 70% ethanol. A total of 50 mL milk was collected by hand into separate tubes and 25 mL was transferred to a sterile 50 mL tube for transport and the other 25 mL was used for compositional Dairy herd improvement (DHI) analysis. For the latter, milk composition (milk fat (%), milk protein (%), lactose (%), solids not fat (%), milk urea nitrogen (%), and SCC (K/mL)) was measured by the Tulare county Dairy Herd Improvement Association using a Fourier Transform Spectrometer 600 Combi System (Combi; Bentley Instruments, Chaska, MN, USA).

Swabs and milk were transported overnight on ice to the Marco laboratory at UC Davis. Upon receipt, the swabs were stored at -20 °C until dislodging the bacteria by incubation in cold phosphate buffered saline (PBS, pH = 7.2; 137 mM NaCl, 2.68 mM KCl, 10.1 mM Na2HPO4, 1.76 mM KH2PO4) on ice with intermittent vortexing every 10 sec for 3 min. Cells were collected by centrifugation at 13,000 g for 10 min at 4 °C and washed twice in PBS before the resulting cell pellets were frozen at -20 °C. For the milk, immediately after receipt at UC Davis, the milk was centrifuged at 13,000 g for 5 min at 4 °C. Cell pellets were then washed twice in PBS before the resulting cell pellets were stored at -20 °C.

### DNA extraction

For teat skin swabs, frozen cell pellets were suspended in Lysis/Binding Solution Concentrate included in MagMax® Total Nucleic acid extraction kit (Thermofisher scientific, Waltham, USA), homogenized with 0.1 mm zirconium beads (Thermo Fisher Scientific, Waltham, USA) using a Fast-Prep®-24 Instrument (MP™ Biomedicals) for 10 sec at 4 m / sec, and then centrifuged for 10 min at 10,000 g. This lysis procedure was previously shown to be effective at retaining an accurate distribution of bacterial taxa in a mock community (26). DNA was then purified using the MagMax® Total Nucleic acid extraction kit (Thermofisher scientific, Waltham, USA) following the manufacturer’s protocol. DNA was stored at -20 °C.

For milk, the protocol used for DNA extraction from swab samples did not result in amplifiable DNA from the freshly expelled milk (data not shown). Therefore, a cetyltrimethylammonium bromide (CTAB) DNA extraction protocol, modified after (27) was used. In short, cell pellets were suspended in 1.2 mL CTAB extraction buffer (100 mM Tris pH 8.0, 1.4 M NaCl, 20 mM EDTA pH 8.0, 5% w/v polyvinylpyrrolidone, 2% w/v CTAB, and 1% v/v beta-mercaptoethanol). The suspension was homogenized in 0.1 mm zirconium beads (Thermo Fisher Scientific, Waltham,WA, USA) in a Fast-Prep®-24 Instrument (MP™ Biomedicals) for 45 sec at 6m/sec and the resulting suspension was centrifuged for 10 min at 10,000 g. Supernatant was collected and incubated at 65 °C for 30 min, with tube inversions every 10 min, prior to centrifugation at 13,500 g for 10 min. Phenol:chloroform:isoamyl alcohol (24:24:1, Thermofisher scientific, Waltham, USA) was added to the lysate and incubated at room temperature for 15 min. This step was repeated and then the upper phase was collected after centrifugation at 13,500 g for 10 min for purification with the PowerSoil® DNA Isolation Kit (MoBio, Carlsbad, CA, USA) according to the manufacturer’s instructions.

### 16S rRNA gene amplicon DNA sequencing and analysis

PCR was carried out using Extaq polymerase (TaKaRa Bio Inc, San Jose, USA) and the 16S rRNA gene V4 hypervariable region (28) 515F (5′-XXXXXXXXCACGGTCGKCGGCGCCATT-3′) and 806R (5’-GGACTACHVGGGTWTCTAAT-3’) primers, including barcodes (random 8-bp barcode) for multiplexing samples attached to the 5’ end of the forward primer (29, 30). PCR was performed under the following conditions: denaturation at 94 °C for 3 min, and 30 cycles of 94 °C for 45 sec, annealing at 55 °C for 60 sec and elongation at 72 °C for 30 sec. PCR amplicons were pooled and eluted from an 1% agarose gel using the Wizard® SV Gel and PCR Clean-Up (Promega, Madison, USA). The Ion Plus Fragment Library Kit (ThermoFisher, Waltham, USA) was used for library preparation and the library quality was assessed using 2100 Bioanalyzer (Agilent, Santa Clara, USA). On-chip library preparation was performed with the Ion Chef™ Instrument (ThermoFisher, Waltham, USA). An Ion Torrent S5 (ThermoFisher, Waltham, USA) was used for sequencing in three different runs. A total of 5,709,613 and 12,760,376 reads were collected for milk and teat skin 16S rRNA V4 amplicons, respectively. A total of 31 milk and 44 skin samples were excluded because they did not yield sufficient reads for further processing and another 24 samples were excluded cause the cow was sold, died or developed mastitis during the study (**Figure S1**).

The demultiplexed, single-end reads were imported into QIIME2 (31). DNA sequences obtained from each sequencing run were processed separately. Reads were denoised by trimming the first 21 bases at the 3’ end and by truncating them at 290 bp. Amplicon Sequence Variants (ASVs) were identified using DADA2(32). A feature classifier based on the SILVA v.138 V3-V4 regions database (33) using the Naive Bayes method was applied for taxonomic classification. Downstream analyses were carried out in R v. 4.1.3 (34) using Tidytacos (35) for visualization and using vegan package (36) for statistics. Inverse Simpson and Observed indexes were calculated for alpha-diversity, whereas the Bray Curtis metric was used to calculate the distance matrices and plot the Principal Coordinate Analysis (PCoA) for beta-diversity.

### Statistical analysis

Due to the sparsity and non-normality of the sample distribution, the non-parametric Wilcoxon test was used for pairwise comparisons between treatments, including Benjamini-Hochberg multiple testing correction. For beta-diversity analysis, Permutational Multivariate Analysis of Variance (PERMANOVA) was used to test impact of environmental factors and metadata on the milk and skin microbiome. This test was performed using the adonis function within the vegan package (37). Results were deemed significant if an adjusted p-value <0.05 was obtained.

## Results

### Milk and teat skin have distinct microbiota

To determine whether the milk and teat skin microbiota were sufficiently similar that the impacts of antibiotic use could be measured for the two sample types combined, the milk and teat skin microbiota collected at baseline (before antibiotic use) were compared. Freshly expelled milk collected at baseline (dry-off) was diverse with on average 64 ± 25 families (102 ± 48 genera) in each sample, among which approximately one-third (24 families) comprised an average relative abundance above 1% per sample. Despite this significant intrasample microbial diversity, a core microbiota (defined as a 90% prevalence and an average relative abundance of >1%) consisting of 11 bacterial families was found in the milk (**Table 1**). These families encompassed an average relative abundance 54.93% in individual milk samples at baseline, and among which *Staphylococcaceae* (11.70 ± 18.5 %) *Peptostreptococcaceae* (8.70 ± 6.23 %) *Lachnospiraceae* (8.41 ± 7.96 %) had the highest mean relative abundance (**Figure 2**).

**Figure 2.**
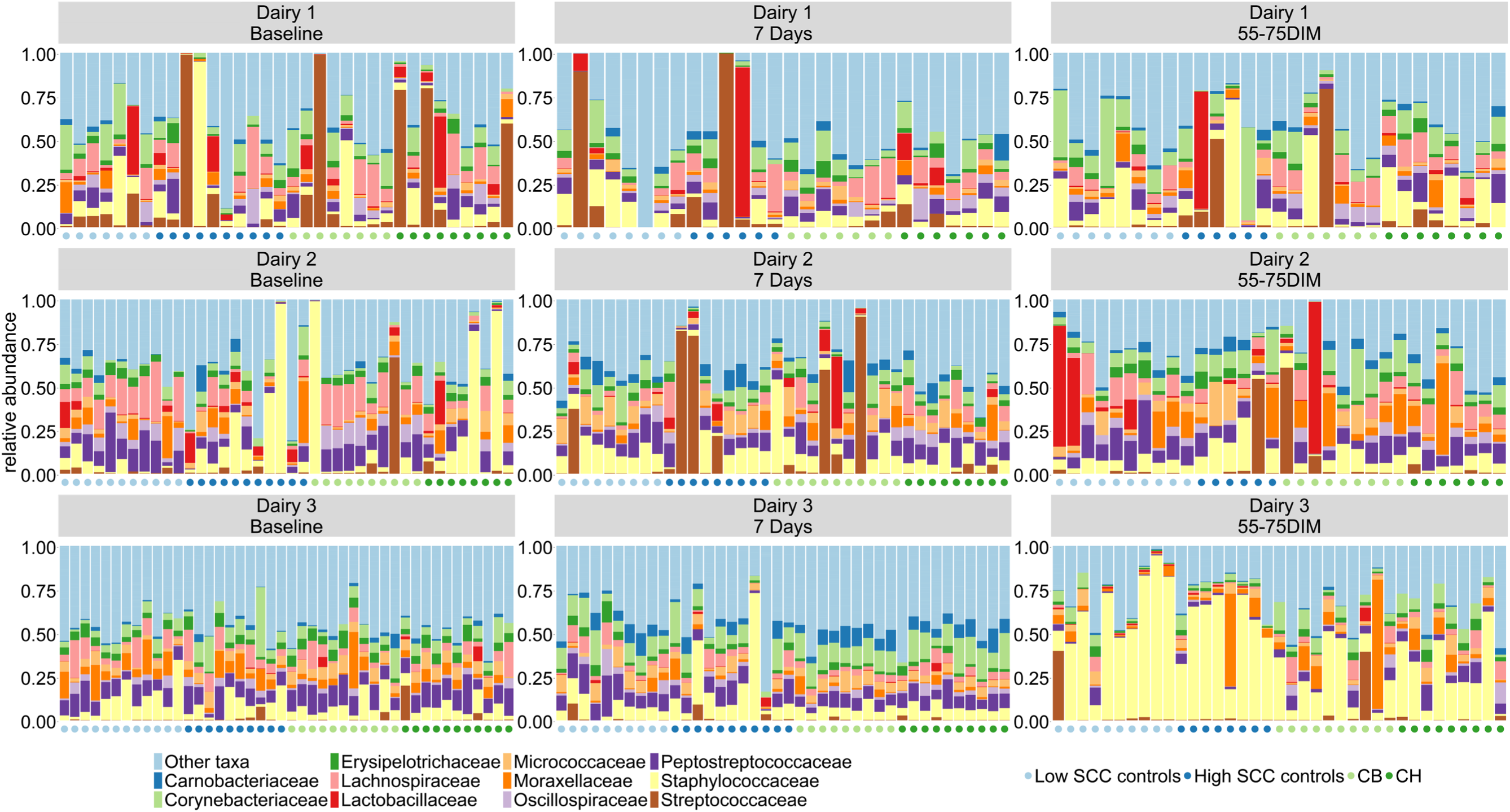
Bacterial taxa in freshly expressed milk across dairies, treatments, and sampling timepoints. The twelve most abundant families are shown. Samples are clustered based on the treatment group of each sample which is indicated by a colored point below each bar. Missing or low read quality samples are not shown.

**Table 1.** Core microbiota^a^ in milk and on the teat skin at dry-off.

| **MILK** |  |  |
| --- | --- | --- |
| **BACTERIAL FAMILY** | **prevalence <sup>b</sup>** | **mean relative abundance (%) <sup>c</sup>** |
| *Staphylococcaceae* | 93.58 | 11.24 |
| *Peptostreptococcaceae* | 100.00 | 8.85 |
| *Lachnospiraceae* | 96.33 | 8.60 |
| *Corynebacteriaceae* | 97.25 | 6.35 |
| *Moraxellaceae* | 99.08 | 4.74 |
| *Erysipelotrichaceae* | 96.33 | 3.54 |
| *Oscillospiraceae* | 95.41 | 3.49 |
| *Micrococcaceae* | 94.50 | 3.33 |
| *Clostridiaceae* | 91.74 | 2.18 |
| *Ruminococcaceae* | 91.74 | 1.42 |
| *Bifidobacteriaceae* | 91.74 | 1.19 |
| **TOTAL** |  | **54.93** |
| **SKIN** |  |  |
| **BACTERIAL FAMILY** | **prevalence** | **mean relative abundance (%)** |
| *Moraxellaceae* | 97.89 | 14.59 |
| *Corynebacteriaceae* | 100.00 | 14.18 |
| *Staphylococcaceae* | 96.84 | 7.43 |
| *Peptostreptococcaceae* | 93.68 | 6.12 |
| *Lachnospiraceae* | 100.00 | 5.07 |
| *Micrococcaceae* | 100.00 | 3.35 |
| *Erysipelotrichaceae* | 90.53 | 2.98 |
| *Oscillospiraceae* | 92.63 | 2.75 |
| *Bacillaceae* | 90.53 | 2.53 |
| *Aerococcaceae* | 94.74 | 2.52 |
| *Bacteroidaceae* | 93.68 | 1.26 |
| *Clostridiaceae* | 94.74 | 1.23 |
| *Streptococcaceae* | 94.74 | 1.17 |
| *Dietziaceae* | 98.95 | 1.11 |
| **TOTAL** |  | **66.29** |

<sup>a</sup> Core microbiota are defined as bacteria present in at least 90% of the milk or teat swab samples at all three dairies and at a relative abundance above 1% at baseline (dry-off).
<sup>b</sup> Prevalence of the bacterial families across all milk and skin samples at baseline
<sup>c</sup> Mean relative abundance of bacterial families per milk and skin sample at baseline

Teat skin had an even more diverse microbiota than milk at baseline, with 84 ± 26 families (49 ± 55 genera) recovered from each skin sample, from which approximately one-fifth (19 families) were present in proportions greater than 1%. The teat skin core microbiota encompassed 14 families comprising an average total proportion of 66.29% per sample (**Table 1**). On the skin *Moraxellaceae* (14.59 ± 16.26 %)*, Corynebacteriaceae* (14.18 ± 13.73 %), *Staphylococcaceae* (7.43 ± 7.45 %) had the highest mean relative abundance (**Figure 3**).

**Figure 3.**
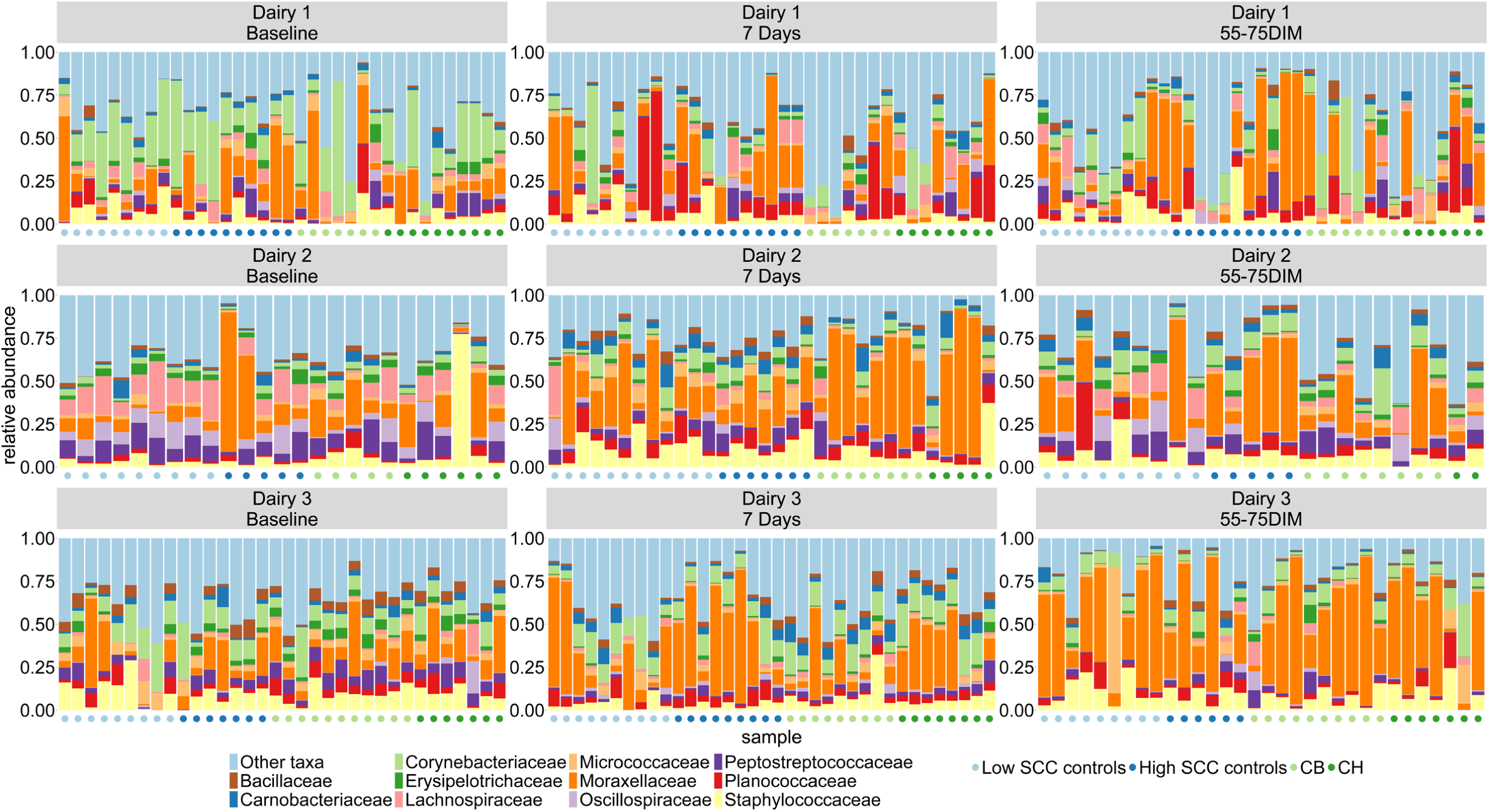
Bacterial composition on teat skin across dairies, treatments, and sampling timepoints. The twelve most abundant families are shown in a stacked bar graph. Samples are clustered based on the treatment group of each sample which is indicated by a colored point below each bar. Missing or low-quality samples are not shown.

Comparisons between the microbiota in milk and on skin at baseline showed that milk had a lower bacterial alpha-diversity compared to skin (**Figure 4A**). Variations between the bacterial contents at the two sites were confirmed by assessment of beta-diversity, showing clear distinction between teat and skin microbiota (Bray-Curtis; ANOSIM, PERMANOVA, p<0.05, **Figure 4B**). These differences were consistent between the three dairies and timepoints (**Figure S2**). At the taxonomic level, nine of the core microbiota families were shared between the milk (9 out of 11) and teat skin (9 out of 14). However, milk contained significantly higher proportions of *Clostridiaceae, Lactobacillaceae, Lachnospiraceae,* and *Oscillospiraceae* and skin had higher proportions of *Corynebacteriaceae* and *Moraxellaceae* (p<0.05, **Figure S3).** At the genus level, milk had significantly higher proportions of *Lactobacillus* and *Staphylococcus* and lower proportions of *Corynebacterium* and *Acinetobacter* compared to the teat skin (p < 0.05,**Figure S4).** Thus, according to the methods applied here, despite having some overlap, the microbiota in freshly expelled milk is distinct from those on the surrounding teat skin.

**Figure 4.**
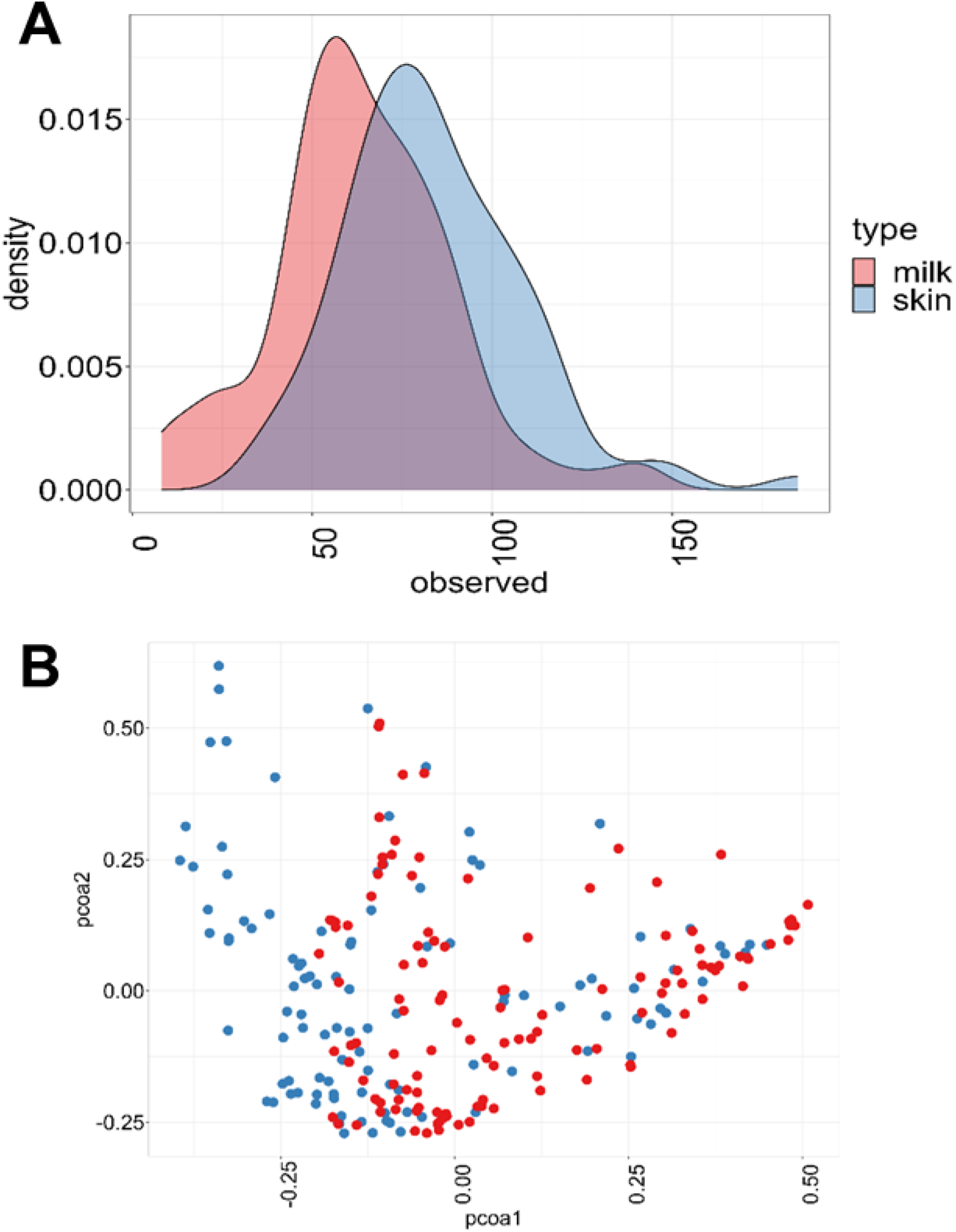
Milk and teat skin harbored a diverse but distinct microbiota at dry-off. **A)** Density plot of the observed alpha diversity of bacterial families in milk and on teat skin at baseline (dry-off). Alpha diversity was calculated as the number of observed families in each milk or teat skin sample at baseline. **B)** PCoA was performed using the Bray-Curtis dissimilarity matrix on ASVs at the family level.

### Correlation between microbial composition and milk composition at baseline

We next assessed whether the milk and teat skin microbiota were correlated with milk composition parameters measured at baseline (dry-off). Small significant correlations were found between SCC and bacterial beta-diversity in milk (PERMANOVA, R^2^= 3.3% p = 0.003), but not with the bacterial beta-diversity on teat skin (PERMANOVA, R^2^= 0.6% p = 0.7). Similarly, SCC was negatively correlated with the inverse Simpson alpha diversity of the bacteria milk (Pearson’s correlation, p=0.02 R^2^ = 13.1%), but not the skin microbiota (Pearson’s correlation, p=0.88, R^2^ = 0.8%). For the other parameters tested (**Table S1**), only Milk Urea Nitrogen (MUN) content was significantly correlated with bacterial beta-diversity in milk (PERMANOVA, R^2^= 12.96% p = 0.019) and with bacterial beta-diversity in teat skin (PERMANOVA, R^2^= 13.31% p = 0.008). In milk, MUN was significantly positively correlated with *Intrasporangiaceae (*R^2^= 31.3%), and *Brevibacteriaceae* (R^2^= 18.2%) whereas *Pseudomonadaceae (*R^2^= -31.3%), and *Streptococcaceae (*R^2^= -11.3%) were negatively correlated with MUN (p<0.05, Pearson’s correlation). On teat skin MUN was significantly positively correlated with *Brevibacteriaceae* (R^2^=24.3%) and *Intrasporangiaceae (*R^2^= 19.7%) and negatively corelated with *Pseudomonadaceae* (R^2^= -24.0%).

### Milk and skin microbiota differ between dairies and timepoints

Because our results strongly indicated that the microbiota detected in freshly expelled milk and on teat skin were significantly different, these sites were examined separately in subsequent analyses. Taking dairy, sampling timepoint, SCC, and antibiotic treatment into account, showed that the main experimental factors explaining the differences in bacterial beta-diversity were the dairy (R^2^ = 6.22 for milk, R^2^ = 7.56 for skin) and time of sampling (R^2^ = 4.74 for milk and R^2^ = 3.77 for skin) (**Table 2**). When examined collectively across timepoints (without taking dairy into account), CB or CH use was associated with a small but significant impact on the milk microbiota diversity (Bray-Curtis, Adonis, p =0.003, R^2^= 1.4%). This difference with antibiotic use was not found for the skin (Bray-Curtis, Adonis, p >0.05, R^2^= 0.8%, **Table 2**). Remarkably, the effect size of SCC was small and not significant (**Table 2**).

**Table 2.** Effect sizes of dairy, timepoint, treatment, and SCC on milk and skin bacterial beta-diversity.

| **Milk** |  |  |  | **Skin** |  |  |
| --- | --- | --- | --- | --- | --- | --- |
|  | **R<sup>2</sup><sup>a</sup>** | **F** | **P-value<sup>b</sup>** | **R<sup>2</sup>** | **F** | **P-value** |
| **Dairy** | 6.22 | 10.98 | 0.001 | 7.56 | 13.37 | 0.001 |
| **Timepoint** | 4.74 | 8.38 | 0.001 | 3.77 | 6.67 | 0.001 |
| **Treatment** | 1.41 | 1.67 | 0.025 | 0.80 | 0.94 | 0.550 |
| **SCC** | 0.42 | 1.51 | 0.129 | 0.28 | 0.98 | 0.423 |

<sup>a</sup> Effect size is shown as the R<sup>2</sup> - value in %.
<sup>b</sup> Impact of experimental factors on beta diversity was calculated using PERMANOVA with the adonis function in the vegan package. 999 permutations were performed.

Further comparisons of the three dairies showed that the bacterial alpha-diversity (as determined by the Inverse Simpson metric) in milk at Dairy 3 was significantly higher than at Dairy 1 and Dairy 2 at baseline, higher than at Dairy 2 at the day 7 sampling timepoint, and significantly lower compared to the other two dairies at 55-75 DIM (**Figure S5A**). For the skin, the bacterial composition on the teat skin at Dairy 1 had the lowest alpha diversity at baseline (p<0.05; **Figure S5B**), however at the day 7 and 55-75 DIM timepoints no significant differences between dairies were observed.

Dairy and timepoint dependent differences in milk microbiota composition were also evident at the taxonomic level (**Figure 3 and Figure S6**). Milk from Dairy 3 had the highest proportion of rare taxa (“other taxa”) at baseline (dry-off) and 7 days later (**Figure S6**). This changed at 55-75 DIM and milk from Dairy 3 was significantly enriched in *Staphylococcaceae* (median relative abundance = 43.3%, min = 1.2% and max = 94.1% for Dairy 3) compared to other two dairies (8.4% and 7.1% median relative abundance for dairy 1 and 2 respectively) (**Figure S6**). Additionally, milk from Dairy 3 contained lower proportions of *Corynebacteriaceae* (median relative abundance = 1.4%, min = 0.04% and max = 14.3% for Dairy 3), *Lachnospiraceae* (median relative abundance = 5.3%, min = 0.2% and max = 16.2% for Dairy 3), and *Peptostreptococcaceae* (median relative abundance = 3.0%, min = 0.2% and max = 14.8% for Dairy 3) than the other two dairies at 55-75 DIM (p < 0.05, Wilcox test) (**Figure S6**). Dairy 1 had significantly lower proportions of *Peptostreptococcaceae* at baseline compared to other two dairies (median relative abundance = 3.6%, min = 0.1% and max = 20.0% for Dairy 1, median relative abundance = 8.7%, min = 0.4% and max = 34.1% for Dairy 2, median relative abundance = 10.1%, min = 1.9% and max = 22.3% for Dairy 3). At 7 days Dairy 1 also had significantly lower proportions of *Peptostreptococcaceae* compared to the other two dairies (median relative abundance = 5.0%, min = 0.2% and max = 12.5% for Dairy 1, median relative abundance = 8.1%, min = 0.3% and max = 18.4% for Dairy 2, median relative abundance = 8.4%, min = 0.8% and max = 23.5% for Dairy 3). Finally, *Streptococcaceae* was enriched in Dairy 1 compared to other two dairies at Baseline and 7 days (**Figure S6**).

For teat skin, there were also significant differences in the proportions of several taxa between dairies and across timepoints (**Figure 4 and Figure S7**). Levels of *Corynebaceteriaceae* were distinct between in the teat skin of all three dairies at baseline (median relative abundance = 22.3%, min = 1.5% and max = 78.8% for Dairy 1, median relative abundance = 3.2%, min = 0.8% and max = 8.8% for Dairy 2, median relative abundance = 11.2%, min = 2.4% and max = 100% for Dairy 3) (**Figure S7**). Significant differences in the proportions of this family were also observed 7 days later between the three dairies.

Additionally, at that timepoint, the proportions of *Peptostreptococcaceae* were significantly higher on the teat skin at Dairy 2 compared to Dairy 3 (median relative abundance = 9.6%, min = 0.6% and max = 21.9% for Dairy 2, median relative abundance = 5.8%, min = 0.3% and max = 28.1% for Dairy 3). (P<0.05) (**Figure S7**). Finally, significant differences in *Staphylococcaceae* relative abundances were observed for teat skin at each timepoint. At baseline, teat skin samples of Dairy 3 (median relative abundance = 9.8%, min = 0.7% and max = 29.0%) had a significantly higher relative abundance of *Staphylococcaceae* compared to Dairy 1 (median relative abundance = 4.2%, min = 0.1% and max = 21.9%) and Dairy 2 (median relative abundance = 3.5%, min = 1.2% and max = 77.4%). After 7 days, Dairy 2 had the highest relative abundance of *Staphylococcaceae* (median relative abundance = 8.0%, min = 1.2% and max = 37.3%), and at 55-75 DIM Dairy 1 had the lowest relative abundance of *Staphylococcaceae* (median relative abundance = 6.8%, min = 0.1% and max = 33.3%) (**Figure S7**).

### Long term effects of antibiotic use were minor and limited to individual dairies and timepoints

Due to the small effect size of antibiotic use on the microbiota in milk (1.4%) and on the skin (1%) and the significant differences between dairies, the impacts of CB and CH were separately assessed for each dairy. Compared to baseline at Dairy 1 and Dairy 2, there was no significant change in bacterial alpha-diversity in milk collected from cows given CB or CH at either day 7 (CB, p=0.6 and CH, p=0.4, Inverse Simpson, Dunn’s test) or 55-75-DIM (CB, p=0.2 and CH, p=0.7). At Dairy 3, bacterial alpha diversity at baseline was significantly higher compared to 55-75 DIM (CB, p=0.03 and CH, p=0.01). However, this effect could not be attributed to antibiotic treatment because the reduction in bacterial diversity was also observed for the controls (low SCC controls and high SCC controls) at that dairy. Moreover, there was no change in the alpha-diversity of bacteria on the teat skin with antibiotic use compared to baseline samples.

To take inter-individual variations in milk and skin microbiota into account, the bacterial composition was compared for each cow over time (baseline and 55-75 DIM) following antibiotic use (**Figure 5**). The Day 7 sampling time point was excluded because at this stage of the lactation cycle the udder is undergoing repair and recovery which leads to high SCC counts. For skin, no significant differences were observed. For Dairy 1 and 2, no significant changes in the beta-diversity of bacteria in milk were found according to this analysis, irrespective of antibiotic treatment (**Figure 5**). For Dairy 3, at 55-75 DIM, milk collected from cows that received CB or CH had a smaller change in bacterial beta-diversity relative to baseline samples compared to the low and high SCC controls **(Figure 5**). This could potentially be linked to the effect of antibiotics on *Staphylococcaceae*, a family which was found in high levels (median relative abundance = 43.3%, min = 1.2% and max = 94.1%) in milk from Dairy 3 at 55-75 DIM across all treatment groups. CB and CH use was associated with small but significant (p< 0.05) reductions in proportions of *Staphylococcaceae* in milk at 55-75 DIM compared to untreated cows (**Figure 6**).

**Figure 5.**
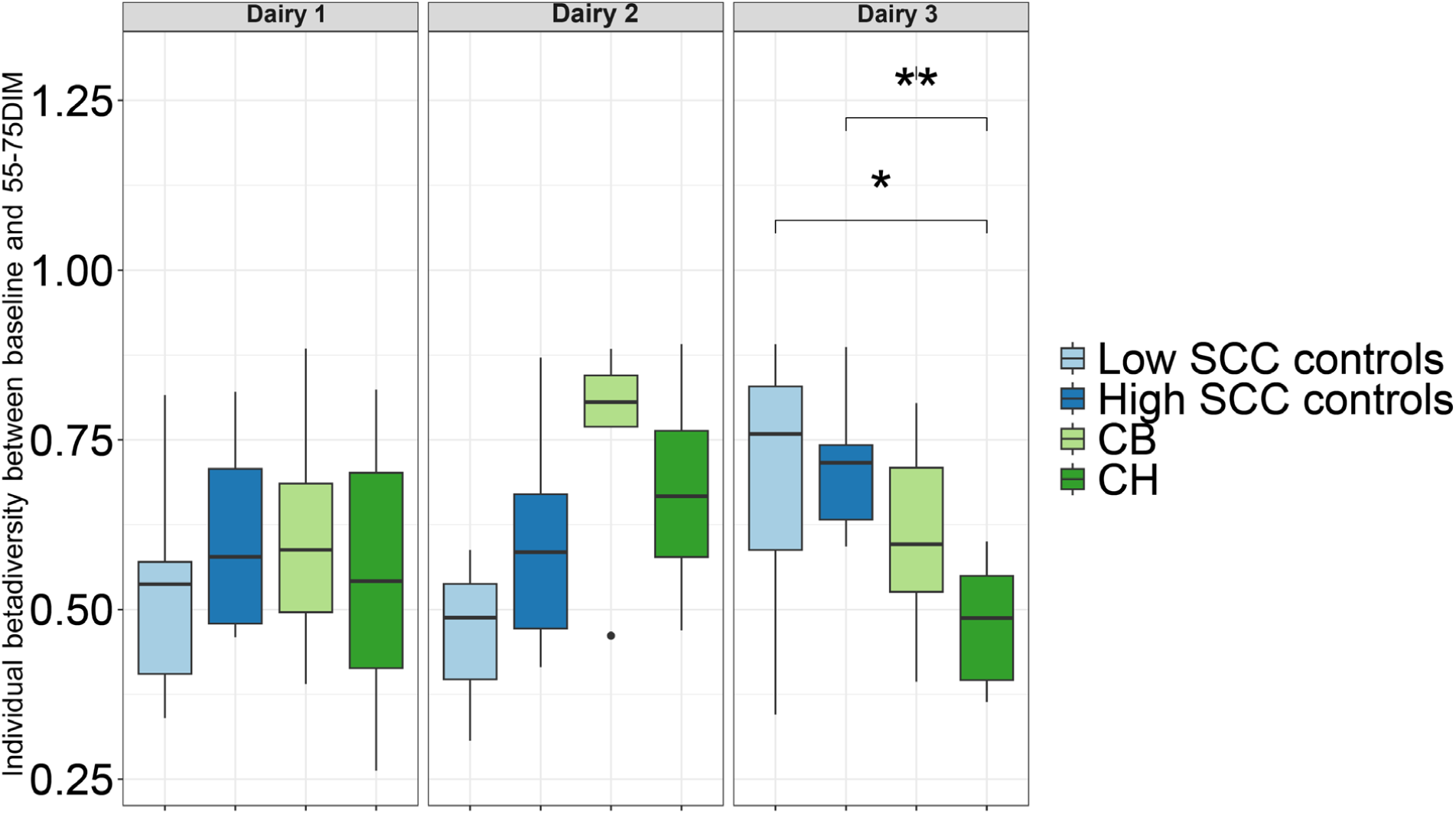
Intra-individual changes in bacterial beta-diversity between baseline and 55-75 DIM. Beta diversity was calculated between milk collected at baseline and 55-75DIM. A higher beta diversity indicates a shift in microbiota composition, with 1 indicating completely different and 0 indicating a completely identical microbiota composition. Significant differences were calculated using a pairwise Wilcoxon test with Benjamini-Hochberg multiple testing correction (p < 0.01 **, p < 0.05 *).

**Figure 6.**
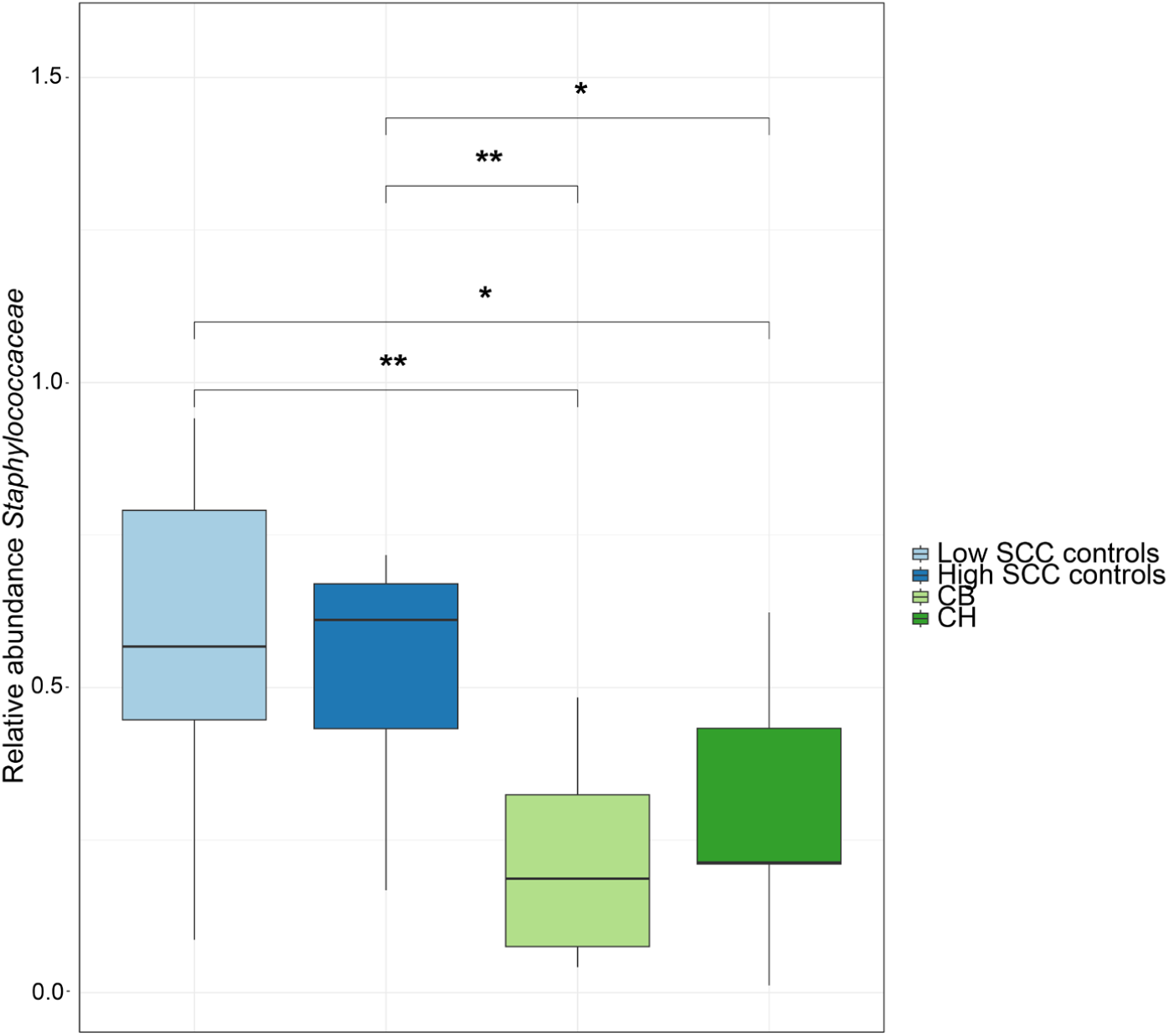
Reduced proportions of *Staphylococcaceae* following antibiotic use at Dairy 3. The relative abundance of *Staphylococca*ceae for each milk sample of Dairy 3 at 55-75 DIM is shown in a boxplot per group. Each boxplot displays median (horizontal line), 25% quantile and 75% quantile in the box. Significant differences were determined using the pairwise Wilcoxon test with Benjamini-Hochberg multiple testing correction (p < 0.01 **, p< 0.05 *).

## Discussion

Understanding the impacts of intramammary antibiotics on the milk and teat microbiota at dry-off is essential for developing improved husbandry practices. These practices should ultimately aim to control mammary infections while minimizing the risk of antibiotic resistance and undesirable perturbations to the mammary microbiota. In this multi-site, time course study, we found that neither of the commonly used cephalosporin antibiotics (CB or CH) resulted in extensive and repeatable changes to milk or skin microbiota on cows at dry-off that had a high SCC but were otherwise healthy without observable mastitis. Only *Staphylococcaceae* levels in milk were affected with antibiotic use according to findings from one dairy and a single time point. The results point to the importance of the environment and time of sampling and potentially the stage of milking on the diversity and proportions of bacteria present in milk and on the udder skin, and thus the need to take these factors into account when planning for microbiome interventions. This study also suggests that using antibiotics at dry-off provides no significant long-term benefits nor harm to cows with high SCC but are otherwise healthy.

Contrary to our understanding of antibiotic use on the gut microbiome (38, 39), intramammary injection of CB or CH did not result in short (7 day) or long-term effects (55-75 DIM into the next lactation) on the milk or teat skin microbiota. This finding is consistent with a prior small-scale study on five healthy cows with low somatic cell counts (SCC) which found no significant effects of antibiotic treatments (cephalonium dihydrate and benzathine cloxacillin) on either the alpha-or beta-diversity of the milk microbiota at dry-off, in the colostrum, or at five days postpartum (40). Similarly, another study with 36 non-mastitic cows housed in the same facility and administered ceftiofur hydrochloride (500 mg; intramammary administration) during the dry-off period found no effect on the milk microbiota or bacterial load at seven days postpartum, nor was there an effect on the SCC score or mastitis incidence (41). Furthermore, in cases of clinical mastitis, no difference in pathogen bacterial load was found between 40 cows given an intramammary injection of ceftiofur hydrochloride (42). These results collectively indicate that non-selective antibiotic therapy does not significantly alter the milk microbiota composition.

The caveat to this main outcome is that antibiotic use was associated with limited effects on certain bacterial taxa at Dairy 3. Cephalosporin treatment may have impacted *Staphylococaceae* at one dairy (Dairy 3), such that in the next lactation cycle (55-75 DIM)a lower relative abundance of *Staphylococcaceae* was observed in the antibiotic treated cows.

This dairy was different from the other two as a large increase of *Staphyloccocacae* in milk was observed, irrespective of SCC and only minor impacted by antibiotic treatment. Thus, cows at Dairy 3 may have had a microbiome more responsive to antibiotic use, indicating that use of targeted antibiotic treatment of ‘at-risk’ cows with higher proportions of specific taxa could be preferred over an untargeted approach.

Broadly, to reach this assessment of antibiotic use on the milk and teat skin microbiota, it was necessary to have a global view on the bacterial diversity at each dairy and the timepoints measured. Both milk and teat skin contained many taxa at a relative abundance below 1%. This observation aligns with previous studies assessing bacterial composition in raw milk (43, 44), in bulk tank milk (45) and on the cow teat skin (11, 46). Despite this diversity, a core microbiota comprised of 11 families was found at all dairies and accounted for approximately 55% of the total bacteria present in each milk sample at baseline. The taxa, predominated by *Staphylococcaceae, Peptostreptococcaceae, Lachnospiraceae,* were also identified as core members of the milk microbiota in other studies (44, 45, 47). A core microbiota of 14 families was also identified for the teat skin and accounted for approximately 66% of the total bacteria in those samples at baseline. The taxa, predominated by *Moraxellaceae, Corynebacteriaceae*, *Staphylococcaceae,* were previously also described as members of the core microbiota on the cow udder (48–50).

Comparisons between the milk and skin microbiota showed that they had many of the same taxa in common. This finding is consistent with prior observations reporting that 70% of bacterial genera are shared between the teat skin and milk of the same cow (17). Interestingly, several of the same bacterial families in milk and on teat skin were significantly correlated with milk urea nitrogen (MUN) levels. High MUN values indicate an excess of nitrogen in the feed that is not utilized by the microbiota and excreted in body fluids. For example, a positive correlation between MUN and *Streptococcus* and other mastitis pathogens (51) indicates a reduction in normal intestinal function. By comparison, negative correlations were found between MUN and the proportions *Pseudomondaceae* in milk and on skin, potentially suggesting that members of these families are associated with a healthy udder. However, similarities between milk and the teat skin microbiota did not extend further. The microbiota in milk may come through various sources, including the skin and potentially other sites such as the canal and apex passage (52) (53, 54). The microbiota on the teat skin can originate from the dairy environment, bedding, among other locations (55–57). Thus, the two sites should be considered to harbor different resident bacterial populations and that the teat skin is not the only source of the bacteria detected in milk.

Dairy farm, time of sampling, but not antibiotic use, had the greatest effect sizes on the milk and skin microbiota. Previous studies conducted in Ireland (44, 58), Italy (59), China (60), and the United States (45), indicated that environmental factors such as geography, altitude, and seasonality significantly influence the microbiota of raw milk. In addition to geographical location, farm-specific conditions such as housing, bedding, and management practices also impact the milk and skin microbiota (50, 61–63). Climate factors like weather and temperature can further influence the milk and skin microbiota at different sampling times from the same location (6, 60).

Although we anticipate our findings are robust, there are a couple limitations to our study. Firstly, we had to use different DNA extraction methods for analysis of the milk and skin microbiota. While it was possible to obtain DNA amenable to PCR from the skin swabs using routine (column based) extraction methods, this method was unsuccessful for raw milk despite repeated attempts, different DNA polymerases and DNA extraction kits (data not shown). Only with incubation of milk in cetrimonium bromide (CTAB) prior to DNA extraction and use of phenol and chloroform (64) was possible to obtain amplification by PCR. CTAB is a cationic detergent that facilitates removing polysaccharides and polyphenols during DNA extraction and has been successfully employed in DNA extraction from milk (65–67) and other complex food matrices, such as olives (27). The necessity to use this more laborious approach on the freshly expressed milk was remarkable since it was not necessary in our prior studies in which raw milk was examined after cooled transport and storage (26, 68). Notably, although these findings our consistent with prior studies (63, 69), it is possible that some of the some of the taxonomic differences observed between these two sites may have been the result of these distinct methodological approaches used. Additionally, because of the highly diverse and low proportions of many taxa, we may have missed changes to taxa that were less abundant and variable between samples. Moreover, due to the use of the short V4-region of the 16S rRNA, our taxonomic resolution is limited. The use of long read, full 16S rRNA amplicon sequencing or shot-gun metagenomic sequencing methods that reduce quantities of host (bovine) DNA (70) could be a good alternative for future experiments to increase the resolution to species level. To assess the effect of antibiotic treated groups, an additional sample for the short-term effects would be beneficial to include in follow up studies.

In conclusion, milk and teat skin harbor a distinct, highly diverse microbiota which is influenced by location and time of sampling during the lactation cycle. Heterogeneity within location microbiota influenced the efficacy of intramammary antibiotic treatment, as no long-term global effects across three dairies of antibiotic treatment was observed. However, limited location and timepoint dependent effects were observed for antibiotic treatment. Thus, while a core microbiota is present, the influence of specific environmental and management factors, along with targeted antibiotic strategies, remains critical in shaping the microbiota and managing mastitis.

## Data availability

Raw sequencing data was uploaded on the European nucleotide archive sequence read archive (ENA-SRA) and is available under the following accession number PRJEB63336.

## Acknowledgements

The research performed was funded by the California Dairy research foundation under grant number: P-21-007-UCD-MM-SR. WVB was partially supported by the Research Foundation – Flanders (FWO, grant 1224923N). We would like to thank Chung Lung Cui and Kainoa Sastra for their help in organizing, processing the milk and swab samples and DNA extractions. We would also like to thank the three dairies in California for their collaboration.

